# Routine Determination of Ice Thickness for Cryo-EM Grids

**DOI:** 10.1101/302018

**Authors:** William J. Rice, Anchi Cheng, Alex J. Noble, Edward T. Eng, Laura Y. Kim, Bridget Carragher, Clinton S. Potter

**Affiliations:** National Resource for Automated Molecular Microscopy, Simons Electron Microscopy Center, New York Structural Biology Center, New York, United States; Department of Biochemistry and Molecular Biophysics, Columbia University, New York, NY 10032, USA

**Keywords:** Cryo-EM, energy filter, mean free path, ice thickness, inelastic scattering

## Abstract

Recent advances in instrumentation and automation have made cryo-EM a popular method for producing near-atomic resolution structures of a variety of proteins and complexes. Sample preparation is still a limiting factor in collecting high quality data. Thickness of the vitreous ice in which the particles are embedded is one of the many variables that need to be optimized for collection of the highest quality data. Here we present two methods, using either an energy filter or scattering outside the objective aperture, to measure ice thickness for potentially every image collected. Unlike geometrical or tomographic methods, these can be implemented directly in the single particle collection workflow without interrupting or significantly slowing down data collection. We describe the methods as implemented into the Leginon/Appion data collection workflow, along with some examples from test cases. Routine monitoring of ice thickness should prove helpful for optimizing sample preparation, data collection, and data processing.

## Introduction

Recent advances in instrumentation have made calculation of near-atomic resolution structures by electron cryo-electron microscopy (cryo-EM) almost routine (Earl et al., 2017; Murata and Wolf, 2018). Detector improvements now give us the ability to correct for image drift, dose-compensate individual frames, and count electrons directly, greatly improving the DQE of the final images, to the point that information can be collected to nearly the Nyquist limit (Ripstein and Rubinstein, 2016). Modern electron microscopes have improved stability, such as constant power lenses, so they can remain well aligned over a several day data collection period (Bierhoff, 2005). Samples can be kept cold for days at a time through the use of autofillers, removing the requirement to fill a dewar every few hours. Various automation programs, such as Leginon (Suloway et al., 2005), SerialEM (Mastronarde, 2005), UCSFImage (Li et al., 2015), and EPU (Thermo Fisher Scientific), have been in place for over a decade, and after initial setup and screening, allow for data collection to proceed over days at a time, providing particle numbers on the order of 10^5^ to 10^6^ from a single grid. These technological improvements, first available on the highest end microscopes, are now migrating to the microscopes previously thought of as screening instruments (Herzik et al., 2017).

While instrumentation has greatly improved, sample preparation remains a significant bottleneck to the generation of high resolution structures. For some samples it can be challenging to get particles into the ice suspended across the holes in the carbon. The most common way of preparing samples, by blotting followed by plunge freezing into liquid ethane or propane (Adrian et al., 1984), has not changed significantly in over 20 years apart from the development of automated blotting devices (Dandey et al., 2018; Feng et al., 2017; Jain et al., 2012; Razinkov et al., 2016; Wei et al., 2018). The vitrification technique takes some time to master and can produce a wide range of ice thicknesses (Fig. 1) across the grid, the square, and even within individual holes. In addition, it was recently demonstrated that most particles adhere to an air-water interface and if the ice is sufficiently thick this may result in collecting images that contain two layers of particles that will be at different defocus levels and potentially overlap in projection (Noble et al., 2018). The ideal ice layer is usually considered to be just thick enough to support the particle in ice, and so on the order of the size of the particle. For these reasons, the ability to determine ice thickness during screening or during data collection is helpful in assessing overall grid quality, and for determining optimal locations for data collection across the grid, and within each hole.

**Figure 1.**
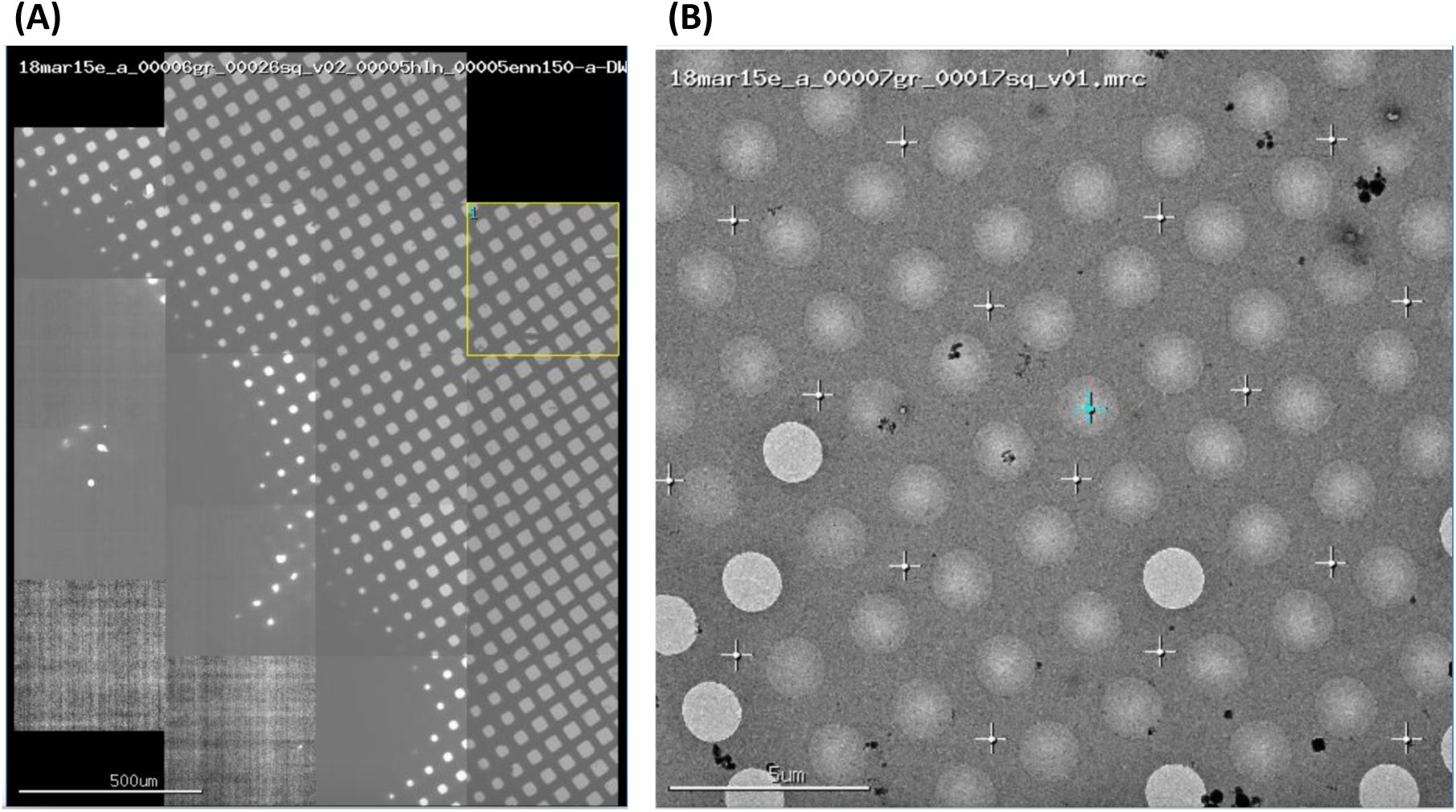
(A) Representative grid atlas collected in Leginon, showing an evident gradient in ice thickness. (B) A “square” level image from this same grid shows evidence of varying thickness, including occasional empty holes.

Ice thickness can be measured in several ways. Possibly the most well defined method is to collect a tomographic tilt series of the desired area, calculate a tomogram, and determine the local ice thickness across the reconstructed area (Noble et al., 2017). This approach is much more time consuming than standard single particle data collection and so is only practical for measuring a few areas, perhaps at the beginning of collection to help determine a targeting strategy. A second method requires tilting the sample to 30 degrees, milling a small hole through the ice, tilting to −30 degrees, and taking an image (Angert et al., 1996). The geometry of this scheme means that the ice thickness can be directly determined by measuring the length of the hole in the second image. While this is easier to do than collecting a full tilt series, it is also disruptive during a data collection run and determining the start and end points of the ice tunnel can be difficult (Feja and Aebi, 1999).

The availability of an energy filter allows direct determination of the ice thickness, either through integration of the energy loss spectrum or through comparison of filtered and unfiltered image intensities (Brydson and Royal Microscopical Society (Great Britain), 2001). This requires knowledge of the inelastic mean free path of the electron through the sample, which depends on voltage, objective aperture width, and sample composition. The relationship is described in equation (1), where d represents the ice thickness, I is the integrated total intensity, I_zlp_ is the integrated zero-loss peak intensity, and Λ is the mean free path for inelastic scattering.

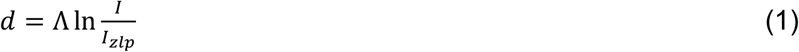

Even in the absence of an energy filter, thickness can be estimated by comparing intensities with and without sample in the electron beam. This calculation depends on the mean free path for elastic scattering outside the objective aperture, which again depends upon voltage, objective aperture width, and sample composition. This is described in equation 2, where d represents the sample thickness, I_0_ is intensity over vacuum, I is the intensity over ice, and λ is the mean free path for elastic scattering outside the aperture.

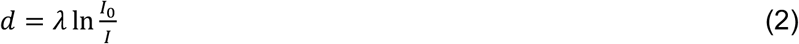

To our knowledge, absolute thickness measurements are not routinely determined during single particle experiments, although the objective scattering method has been described (Yan et al., 2015). Determination through tilting is understandably more difficult and unlikely to be performed as a matter of course. However, estimates from intensity, either with or without an energy filter, can be used to measure the thickness using automated software with minimal additional configuration and time overhead. What is lacking are accurate values for inelastic and elastic mean free paths. These values have previously been determined empirically as well as calculated based on scattering theory, mostly at 80-120 keV, and the published results exhibit a wide variation (see (Vulovic et al., 2013) for a plot of several values). Part of the variance in mean free path for inelastic scattering comes from the fact that some electrons will also be lost due to elastic scattering outside of the objective, and so measurements for inelastic scattering will have an elastic component (Feja and Aebi, 1999; Grimm et al., 1996). Values for both elastic and inelastic mean free paths will therefore vary depending on the acceptance angle for the objective aperture, though for a specific microscope type and objective aperture diameter they should be constant.

We have measured the mean free path for inelastic scattering on an FEI Titan Krios (300keV) equipped with a BioQuantum GIF, and the mean free path for objective elastic scattering on an FEI Titan Krios (300keV), an FEI BioTwin T12 (120 keV), and a Tecnai F20 (200 keV). With these values we have implemented a method into our standard Leginon/Appion data collection workflow (Lander et al., 2009; Suloway et al., 2005) so users can routinely monitor ice thickness. Here we will describe the methodology and the results from a few of our standard test specimens. The methods described in this paper could be readily implemented on other microscopes and automated data collection workflows.

## Methods

### Specimen preparation

Samples were plunge frozen using standard techniques on a Gatan CP3 plunge freezer or a Leica plunge freezer. Rabbit muscle aldolase was prepared and frozen on gold Ultrafoil grids according to (Herzik et al., 2017). Proteasome grids were prepared and frozen on Quantifoil grids according to (Campbell et al., 2015). Glutamate dehydrogenase was prepared and frozen according to (Merk et al., 2016).

### Microscopy

Images were collected on several microscopes and cameras: Titan Krios with energy filter, 100 μm objective aperture, Gatan Bioquantum K2, dose rate 8 e^−^/pix/sec.

Titan Krios with energy filter and Cs corrector, 100 μm objective aperture, Gatan Bioquantum K2, dose rate 8 e^−^/pix/sec.

Titan Krios, Gatan K2, 100 μm objective aperture, dose rate 8 e^−^/pix/sec.

Tecnai F20, 70 or 100 μm objective aperture, DE20 direct detector, dose rate 2 e^−^/pix/frame.

Tecnai T12, 70 or 100 μm objective aperture, TVIPS F416 CMOS detector.

### Tomography

Tilt series were collected on the Titan Krios microscopes using the Tomography app as implemented in Leginon (Suloway et al., 2009). Tilt series were collected on the T12 and F20 microscopes using SerialEM software (Mastronarde, 2005). In both cases, tilt series were collected bidirectionally between −45 and + 45 degrees with a starting angle of 0 degrees and an angular increment of 3 degrees. In all cases, tomograms were calculated using Protomo software as implemented in Appion (Noble and Stagg, 2015).

### Ice thickness measurement from tomograms

Based on the results of hundreds of tomograms (Noble et al., 2017), we know that the vast majority of particles on a vitrified grid are closely associated with the air water interface. We can thus estimate thickness from the z height between proteins (or ice contamination) observed on the two interfaces. For very thin samples the thickness was estimated from this single layer. For thicker samples which had a varying thickness, we chose the approximate average thickness.

### Single particle dataset collection

Images were collected using the Leginon workflow (Suloway et al., 2005). For ice thickness determination by inelastic scattering, two extra 0.5s images were collected; the first image with no slit inserted and the second with a 15 eV slit inserted. For objective scattering ice thickness measurements, several images were taken at the start of the session over vacuum and the mean of these intensities was used as a reference for I_0_.

### Image calculations

Various image calculations and plots were calculated using the EMAN2/Sparx suite of image analysis tools (Hohn et al., 2007; Ludtke, 2016). CTF measurements and Thon ring extent were done using CTFFIND4 (Rohou and Grigorieff, 2015). Curve fitting and plotting was performed using gnuplot, an open source plotting tool.

## Results and Discussion

### Determination of the mean free path for inelastic scattering

The only parameter needed for determining ice thickness when using an energy filter is the mean free path for inelastic scattering in ice. This value will vary depending on the protein concentration, but not significantly. We measured ice thickness from tomograms of two samples (T20s proteasome and rabbit muscle aldolase). We then took pairs of images, with and without the energy filter slit inserted, to get values for I and I_zlp_. A slit width of 15 eV was chosen to eliminate most of the ice and organic plasmon and structure peaks (Leapman and Sun, 1995). A plot of thickness versus ln(I/I_zlp_) is linear, with slope 395 +/− 11 nm (Fig. 2A). This value agrees well with the previously determined value of 400 nm by (Yonekura et al., 2006), on a 300 keV Polara F30 with 70 μm objective aperture. Images of areas of thicker ice occasionally have varying values across the area, which made a single number hard to determine. Nevertheless, since the log ratio of intensities is multiplied directly by this slope value, it should be accurate enough for routine measurement. As a further verification of the method, we acquired pairs of images at 0 degree and 45 degree nominal tilt, and determined thickness using equation (1) and the mean free path as determined above and as expected the thickness increased by a factor of 1.4 (Fig. 2B).

**Figure 2.**
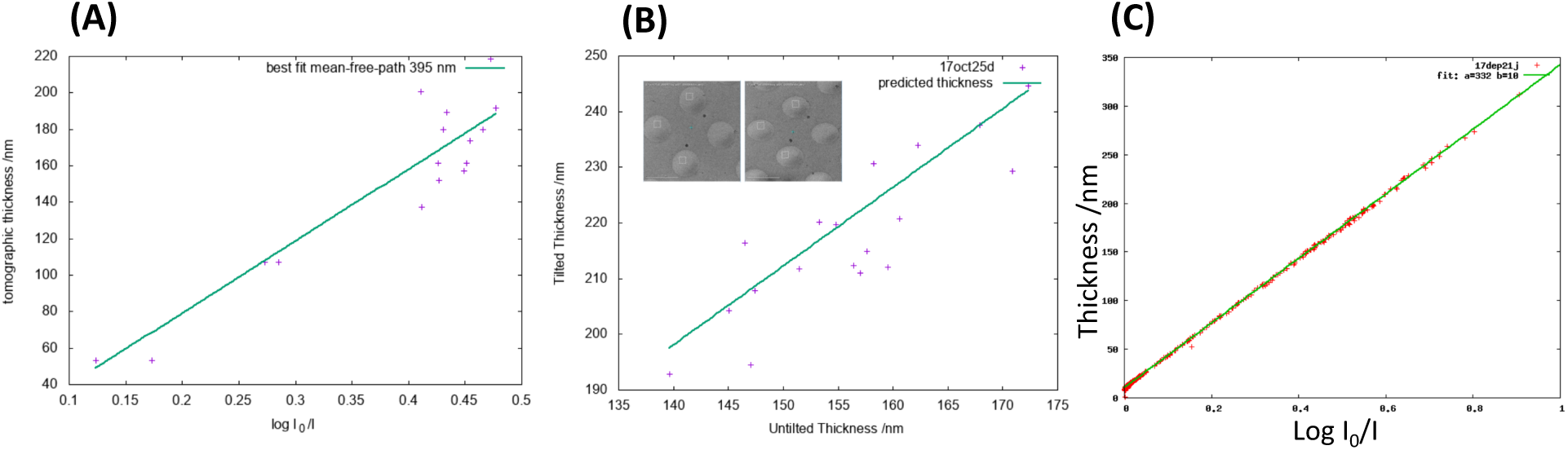
(A) Determination of mean free path for inelastic scattering by electron tomography. Images were collected for both aldolase and proteasome samples with and without a 15 eV energy slit. Tomograms were collected at the same areas as thickness was measured. Thickness versus Log (_Itot_/I_ZLP_) is plotted. The line of best fit (green) had a slope of 395 nm. (B) Thickness determined using the energy filter for untilted and 45 degree tilted images. Inset: an overview of 4 holes are shown where the measurements were made, untilted (left) and tilted (right). For each hole, thickness was measured as described for both tilted and untilted images, and plotted as tilted thickness versus untilted thickness (purple crosses). The green line is a plot of the equation y=1.41x, which is where the points should ideally lie. (C) Determination of elastic mean free path scattering using energy filter thickness. Thickness was measured over many images using the energy filter. Thickness plotted versus log I_0_/I as described in the text. For this experiment, the slope was 332 nm.

### Determination of the mean free path for elastic scattering

The determination of ice thickness using an energy filter is convenient, but this method has two disadvantages: it requires a microscope equipped with an energy filter, and two extra images must be acquired. These images are taken automatically using a very short exposure time, and need not be done for every exposure, but they do add a slight overhead to the collection time. An alternative option is to use elastic scattering outside the objective aperture for this measurement, which can be done on any microscope. In order to use this method, it is first necessary to determine the mean free path of objective scattering, and to collect a vacuum image using the same imaging conditions as used for collection. Application of equation (2) then provides the thickness. The disadvantage of this method is that it relies on the beam intensity being constant throughout the collection, although this can be monitored and a new vacuum image can be acquired when necessary.

In order to determine the value of λ on microscopes with an energy filter, we measured ice thickness using tomography, then used this value in equation 2 to solve for λ. A plot of thickness versus log (I_0_/I) is shown in Fig. 2c. The plot is highly linear, with a slope of 320 nm. We repeated the experiment three times, using both T20S proteasome and aldolase samples, with an average value of 322 nm (sd=6 nm; n=4).

For microscopes without an energy filter, we used several methods to determine the scattering factor for the conditions under study. The first method was to collect tomograms at various locations to calculate thickness, then measure intensities over the same or similar holes and apply equation 2 to determine λ. This was done using both the thin aldolase sample and a thicker proteasome sample. The second method was to use the aldolase sample frozen on gold grids as a ruler to get a defined ice thickness. Our studies of this sample show that we get a very reproducible average ice thickness of 15-20 nm for most images. For each microscope, we collected a reasonably sized dataset (at least 100 images). For each image, we calculated a value of log(I_0_/I), calculated mean and standard deviation values for the set, discarded values more than 2 standard deviations from the mean, and collected a new mean value from the pruned set. Application of equation (2) using the known ice thickness of 15 nm gave us a measure of λ for inelastic scattering outside the aperture. The final method, used for determining scattering factor for a second objective aperture once it was determined for the first, was to measure intensities over the same area using both apertures, calculate the thickness from the first then determine the factor for the second aperture by rearranging equation 2. These results are summarized in Table 1.

**Table 1:**
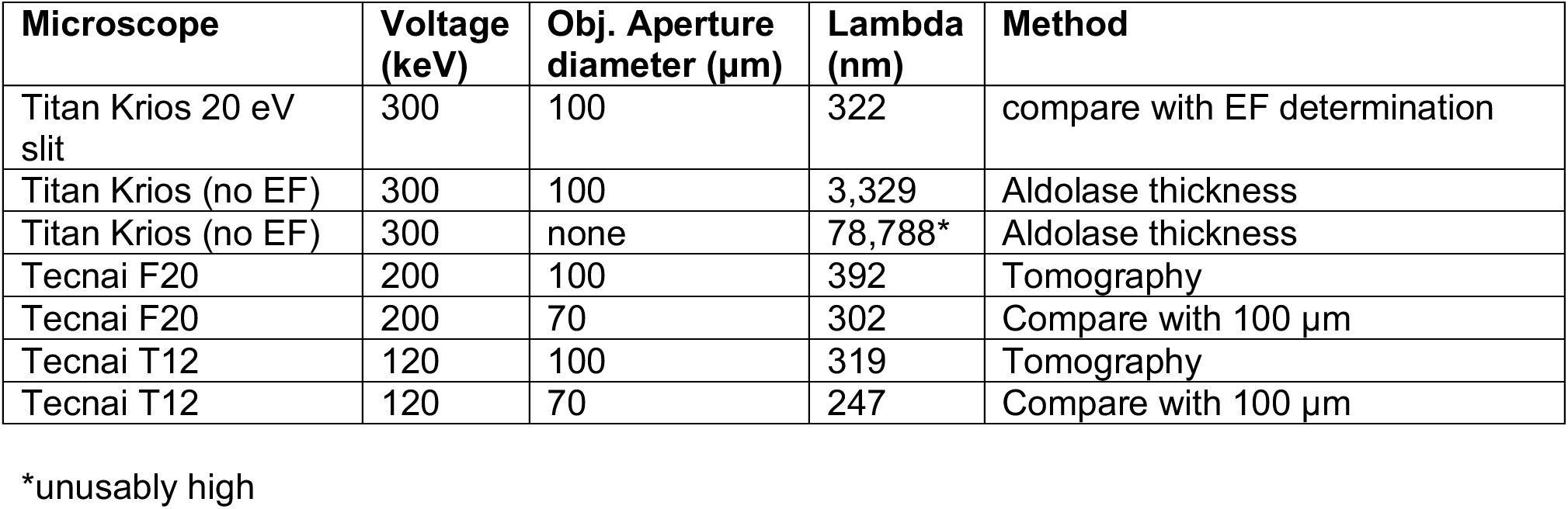
Mean free path for objective scattering.

| Microscope | Voltage (keV) | Obj. Aperture diameter (μm) | Lambda (nm) | Method |
| --- | --- | --- | --- | --- |
| Titan Krios 20 eV slit | 300 | 100 | 322 | compare with EF determination |
| Titan Krios (no EF) | 300 | 100 | 3,329 | Aldolase thickness |
| Titan Krios (no EF) | 300 | none | 78,788* | Aldolase thickness |
| Tecnai F20 | 200 | 100 | 392 | Tomography |
| Tecnai F20 | 200 | 70 | 302 | Compare with 100 μm |
| Tecnai T12 | 120 | 100 | 319 | Tomography |
| Tecnai T12 | 120 | 70 | 247 | Compare with 100 μm |

\*unusably high

These results show that a value of 300 nm is a good starting estimate for elastic scattering MFP on both the T12 and F20 microscopes or the Titan Krios when an energy filter is inserted, despite voltage differences. For the Titan Krios without an energy filter, the value is 10-fold higher, presumably due to the fact that it includes information out to about 1.4 Å resolution. Without an objective aperture inserted on this microscope, there is very little loss by scattering and so the method will not work because of the high value of lambda.

### Application to test samples

As a test specimen, we collected a high quality dataset on our Titan Krios microscope equipped with Cs corrector and Gatan Bioquantum energy filter. This was an exquisitely thin sample, approximately 15 nm on average, and our standard processing pipeline resulted in a structure of 2.4 A resolution (further details in Kim et al, submitted). A histogram of ice thickness for this sample is shown in Fig. 3A. The majority of the images had extremely thin ice: less than 20 nm as measured by energy filtration. Considering the long dimension of the protein is ∼10 nm, the ice was thick enough to just support a single layer of particles. The protein was also very tightly packed together, and perhaps this close packing helps to support an extremely thin layer of ice. The sample was frozen on a gold foil rather than carbon, and gold foils have been shown to result in more consistent ice in general (Russo and Passmore, 2014). We found that, for this sample, the use of carbon grids resulted in either less tightly packed or more aggregated protein and thicker ice overall (Kim et al, manuscript under review).

The proteasome test specimen, in contrast, showed much thicker ice overall and a greater variation in thickness (Fig. 3B). Despite this increased thickness, this dataset also went to high resolution (2.6Å, Kim et al, manuscript under review). Note that the proteasome is larger, at nearly 20 nm for its longest dimension. Tomography by Noble et al. (Noble et al., 2018) showed that most of the sample is at one of the air-water interfaces, which means that most of the particles are at the same height despite the increased ice depth. Additionally, it is approximately 700 kDa in molecular weight, making it easier to see in thicker ice than the 150 kDa aldolase protein.

**Figure 3.**
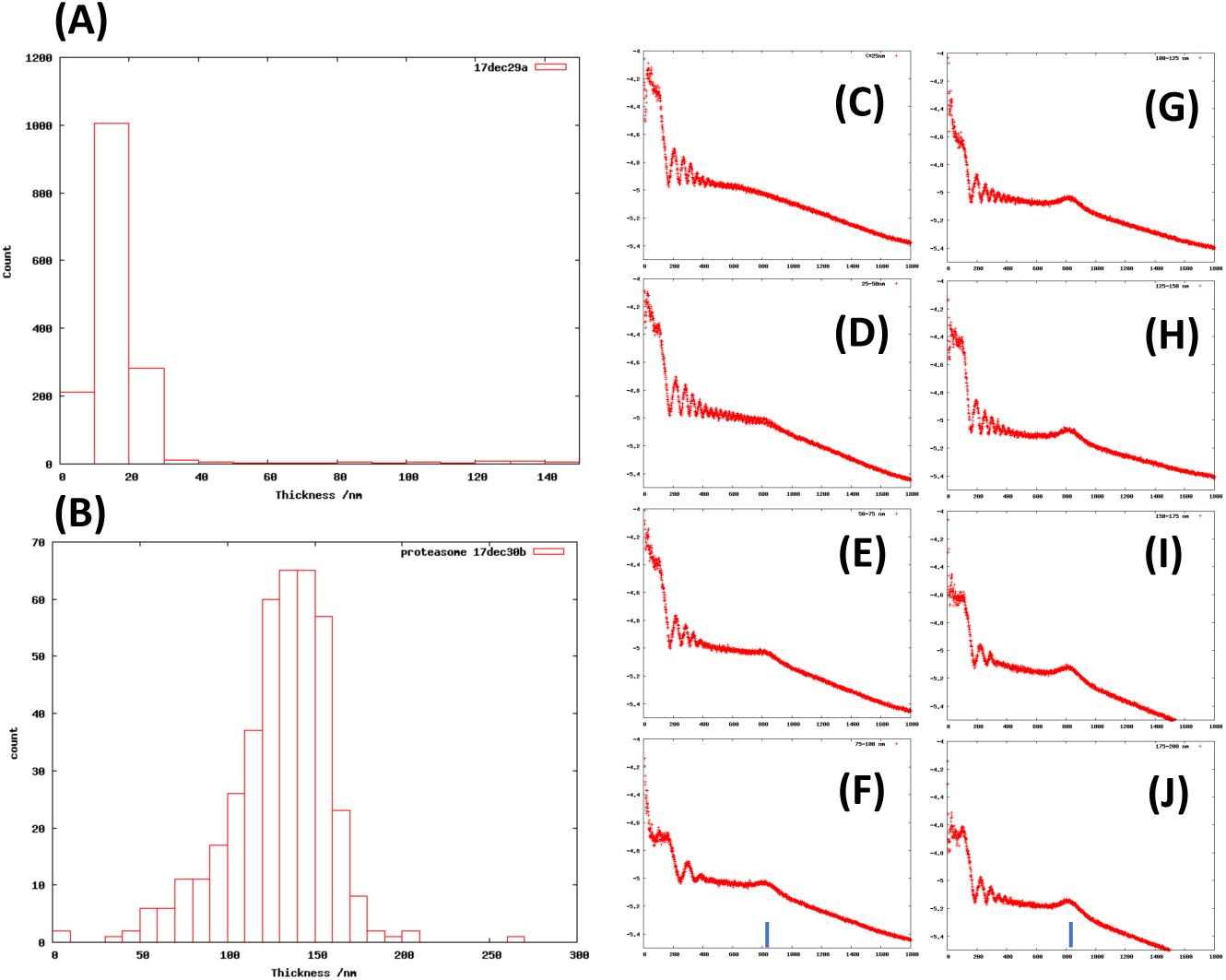
(A) Histogram of thickness values as measured on a rabbit muscle aldolase sample frozen on gold Ultrafoil grids. (C) Histogram of thickness values as measured on a T20S proteasome test sample frozen on C-flat carbon grids. (C-J) Plots of mean radial intensity of Fourier transforms of image averages versus resolution for various ice thicknesses. Blue bar:
3.9 Å^−1^ resolution. (C): 0-25 nm thickness. (D): 25-50 nm thickness. (E): 50-75 nm thickness. (F): 75-100 nm thickness. (G): 100-125 nm thickness. (H): 125-150 nm thickness. (I): 150-175 nm thickness. (J): 175-200 nm thickness.

Sorting the images based on ice thickness revealed some interesting trends. Dividing the micrographs into bins of 25 nm thickness (0-25 nm, 25-50 nm …, 175-200 nm) and plotting the 1D radial averages of summed power spectra showed that the ice ring at 3.9 Å resolution only appears once the ice is 50-75 nm thick, with increasing prominence as the ice layer thickens (Fig. 3 (C)-(J)). Thus, the appearance of this ring gives immediate feedback about the overall thickness of the sample. This sort of information is also collected in the Focus package (Biyani et al., 2017), where it is listed as an “iciness” metadata value for the micrograph. Plotting ice thickness versus Thon ring extent showed that images with the thinnest ice had the highest resolution Thon rings, apart from where the ice was too thin or absent. As ice becomes thicker, resolution steadily worsens. Interestingly, we often notice two separate lines as ice thickens, indicating a subset of images still goes to high resolution in spite of thicker ice (Fig. 4). We have seen this trend on several samples. While we don’t have a definitive explanation, one possibility is that some areas have most particles on one of the air-water interfaces, thus putting most of the scattering at the same height, Other areas of thick ice may have particles on both interfaces and/or scattered throughout the layer, resulting in a less coherent CTF. Examples of both distributions have been observed (Noble et al., 2017).

**Figure 4:**
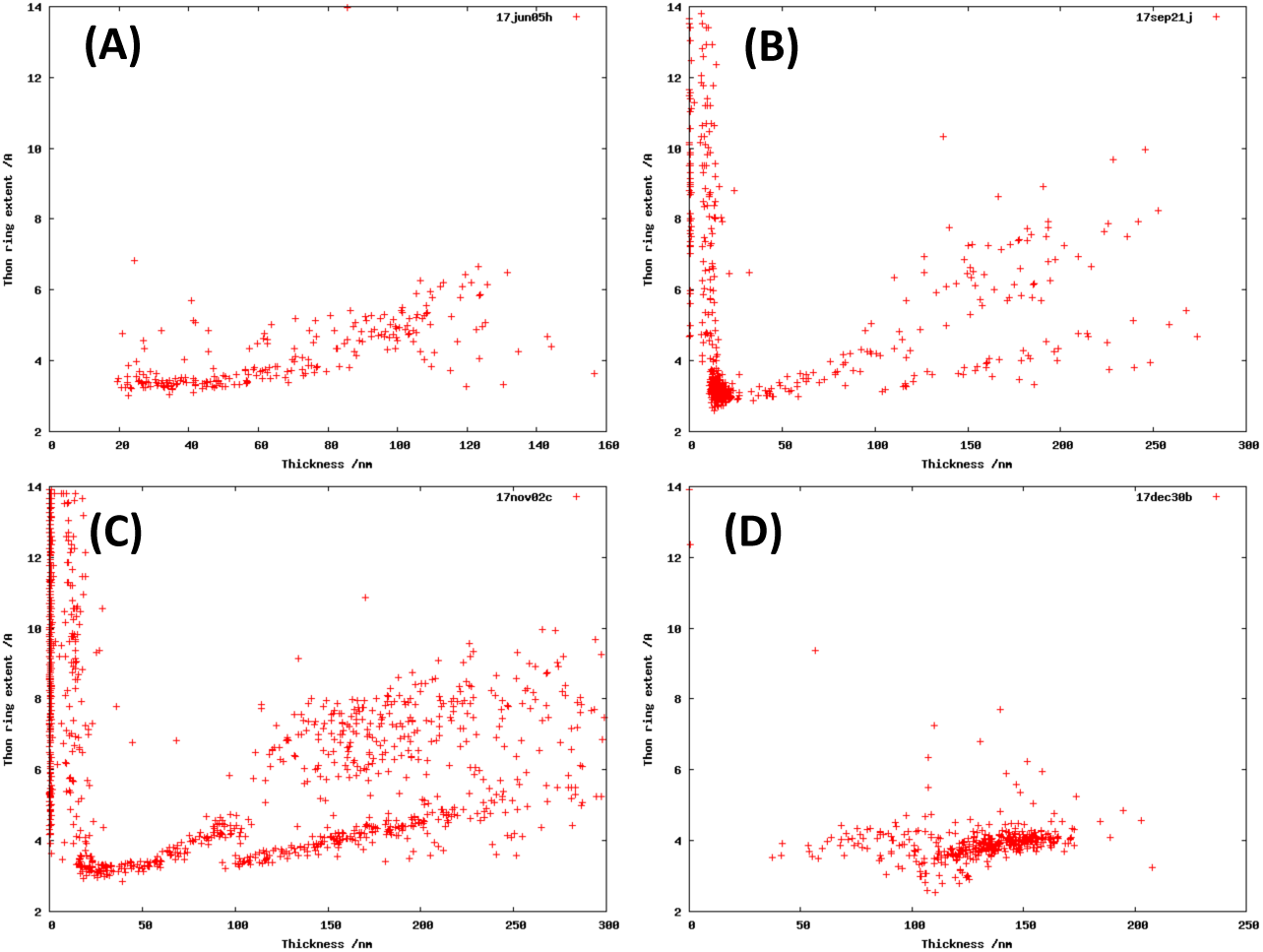
Plots of Thon ring extent, as measured by CTFFIND4 (Å^−1^), versus ice thickness for several samples. (A): glutamate dehydrogenase. (B, C): rabbit muscle aldolase. (D): T20S proteasome. Thon ring extent goes to very low resolution where the ice is too thin or absent, is optimal at thin ice, and generally rises as ice thickens.

### Implementation

We have implemented both methods of ice thickness determination into a new node in Leginon, named iceT (Fig. 5A), and results are shown in the Leginon web interface (Fig. 5B,C). For thickness determination by energy filtration, the user needs to input the mean free path, exposure times, slit width, and how frequently to take the measurement. This feature could be readily added to other collection suites such as SerialEM (Mastronarde, 2005) and EPU. Thickness determination by objective scattering can be done on every image with no loss in time, with the caveat that any change in beam intensity will throw off the calculation. On microscopes with an energy filter, we found that measurement by objective scattering was virtually identical to measurement using the energy filter. Therefore, we now measure thickness with the filter for every image at the start of a session during the initial screen, then reduce to every 10 or 20 images once fully automated collection starts. On high-end microscopes, beam intensity is generally constant throughout a 1 or 2 day imaging session. For screening microscopes, it would be advisable to occasionally check intensity over vacuum and adjust the reference brightness parameter accordingly. We have found that routine monitoring of ice thickness has been useful in both guiding targets and characterizing samples.

**Figure 5:**
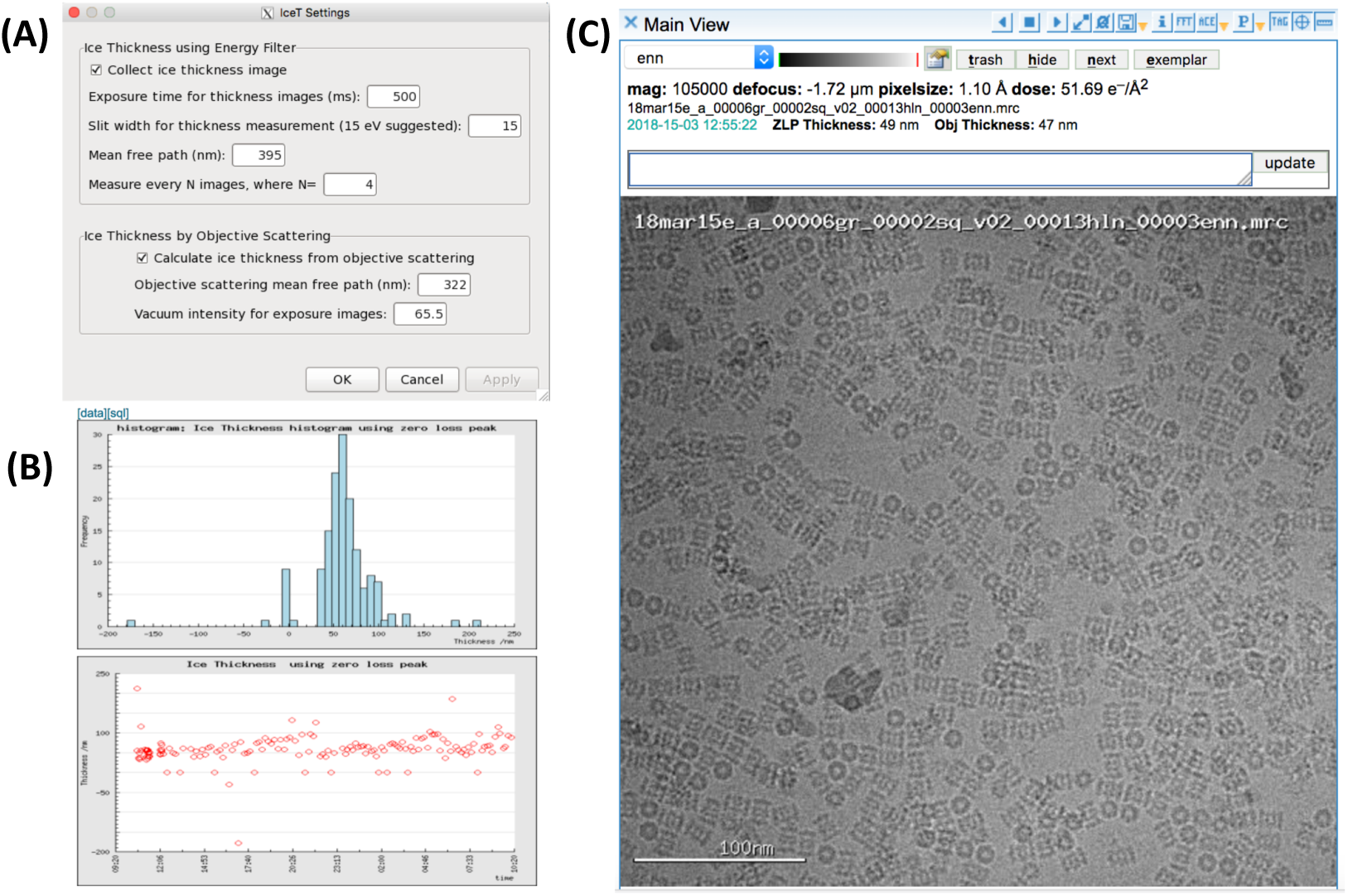
Integration of ice thickness determination into Leginon. (A) The control panel for the node is shown. The user provides various parameters for thickness determination and chooses how often the thickness is measured. (B) Thickness plots for an experiment as shown in the Appion summary pages. (C) Thickness information is displayed when viewing the image in the Leginon web interface.

## Acknowledgements

This work was performed at the Simons Electron Microscopy Center and National Resource for Automated Molecular Microscopy located at the New York Structural Biology Center, supported by grants from the Simons Foundation (349247), NYSTAR, and the NIH National Institute of General Medical Sciences (GM103310) with additional support from the Agouron Institute [Grant Number: F00316] and NIH S10 OD019994-01. We thank Carl Negro (New York Structural Biology Center) for technical assistance with php code. We thank Yifan Cheng (UCSF) for the gift of the T20S proteasome sample.

